# DRUG TARGET IDENTIFICATION VIA A CONDITIONALLY STABILIZED TurboID ENZYME

**DOI:** 10.64898/2026.05.05.722860

**Authors:** Yi Xue, Fynn Zaczek, Ralf-Peter Jansen

## Abstract

Different small-molecule drugs targeting the same protein can produce divergent clinical outcomes through poorly characterized interactome changes. Existing proximity labeling approaches for target identification suffer from background biotinylation independent of small-molecule recruitment, obscuring true drug targets and their binding partners. Here, we incorporate a destabilizing domain (DD) into the biotin targeting chimera (BioTAC) framework to create ddBioTAC, wherein the proximity labeling enzyme TurboID is selectively stabilized only upon binding of a bifunctional targeting molecule. Using the bromodomain-targeting molecule NICE-01 in HeLa cells, we demonstrate that, in the absence of the bifunctional targeting molecule the destabilized TurboID enzyme (TurboID-DD) exhibits reduced protein levels and biotinylation activity compared to the control TurboID-FKBP (FK506-binding protein), while recovering comparable activity upon NICE-01 treatment. This results in an eightfold improvement in specific enrichment of the known target bromodomain containing protein 4 (BRD4) and its interactors, including MED1 and EF1D. Proteome-wide mass spectrometry confirms that ddBioTAC more accurately discriminates drug targets and proximal interactors from non-specific background, advancing unbiased drug-induced interactome profiling.

## Introduction

Small-molecule based therapies are central to modern medicine, yet drugs that bind the same target with similar affinity can produce markedly different clinical outcomes ^[1]^. Such divergent effects can arise from uncharacterized off-target activities or drug-induced changes in the protein interactomes through allosteric or molecular-glue mechanisms ^[2]^. Despite their importance, these effects are poorly characterized for most ligands, creating a major blind spot in drug discovery workflows.

Current unbiased target-identification methods, such as photoaffinity labeling and μMap, rely on light-activated chemistries to tag drug-bound proteins ^[3]^. While powerful for identifying primary targets, their short labeling radius and cell-permeability constraints limit detection to direct binders and preclude mapping of drug-bound protein complexes or *in vivo* applications ^[4]^. In contrast, affinity purification and proximity labeling methods (e.g., BioID, TurboID) can capture broader interactomes in cells and organisms, including transient interactions ^[3a, 5]^. However, these approaches depend on prior knowledge of the target protein and its fusion to the labeling enzyme, which can perturb native interactions and prevent unbiased small-molecule profiling. Furthermore, they can result in extended lists of low-stringency interacting proteins that complicate interpretation of the results.

To enable systematic discovery of drug-induced interactome changes, small-molecule based proximity labeling approaches (e.g. target-ID, PROCID, BioTAC) have been developed that direct the labeling enzyme by a bifunctional molecule to an non-engineered target protein ^[6]^. One of these is the biotin targeting chimera (BioTAC) approach ^[6b]^. Here, the biotin ligase miniTurbo is fused to a mutant version of a FKBP12 domain (FKBP^F36V^), which binds to a recruiter molecule orthoAP1867. By adding a bifunctional molecule assembled from the recruiter and a drug of interest, the biotin ligase can be directed to the drug’s target(s). Proximity labeling can thus result not only in biotinylation of the drug target itself but also of proteins associated with the target of interest ^[6b]^.

BioTAC as well as other enzyme-based proximity labeling approaches suffer from background labeling ^[6a, 6b]^, e.g. caused by the activity of the BioID enzymes prior to its drug-mediated targeting. To control the unwanted or premature activation of proximity labeling enzymes, light-activated variants (Shafraz, 2023; Lee 2023) or APEX2 peroxidase based systems ^[7]^ have been introduced that allow controllable activation of the enzyme.

An alternative strategy to these systems could be a stabilization of only the targeted or bound enzyme while the unbound fraction is degraded. Here, we present ddBioTAC (destruction domain assisted biotin targeting chimera), a proximity labeling system using a destabilized TurboID enzyme that results in improved signal-versus-background labeling. ddBioTAC was benchmarked against the original BioTAC using NICE-01, a bifunctional molecule targeting bromodomain containing proteins like BRD4 ^[8]^. In a direct comparison, ddBioTAC not only leads to significantly higher specific enrichment of the target but also achieves a more reliable identification of BRD4-associated proteins, demonstrating that ddBioTAC represents a useful second-generation proximity labeling system.

## Results and Discussion

### A destabilized TurboID enzyme for targetID

A common issue with target identification(targetID) using an enzyme-based proximity labelling approach is that the biotinylation activity of the proximity ligation (PL) enzyme is independent of the presence of the targeting small molecule. This results in unwanted background biotinylation, which can obscure identification of the correct target or target partners ^[6a, 9]^. Previous studies have demonstrated that the addition of small, membrane-permeable compounds can stabilize proteins fused to this domain, including GFP, Cas9 or Suntag ^[10]^.

To benchmark the effect of a destabilising domain on TurboID in a targetID approach, we compared its stability, biotinylation activity and specificity with those of a related fusion protein that binds the same small molecule, but whose stability is not controlled by binding (Figure 1A and 1B). For the latter, we chose the bifunctional molecule NICE-01 ^[8]^ as the effects of its target recruitment and stabilisation have been well studied ^[11]^. NICE-01 contains the FKBP12 recruiter ortho-AP1867 (Shld1; ^[11]^), which binds to the FKBP^F36V^ and FKBP^F36V L106P^ mutant versions of the FK506-binding protein (Figure 1). The latter is referred to as the DD domain and leads to destabilisation of a protein to which it is fused, but it is stabilised upon binding of Shld1. Shld1 is linked to JQ1 (Figure 1C), a small molecule that selectively targets the bromodomain of the bromodomain and extra-terminal (BET) family ^[12]^. Both fusion proteins are transiently expressed in HeLa cells from a CMV promoter and carry an N-terminal V5 tag for detection (Figure 1D).

**Figure 1.**
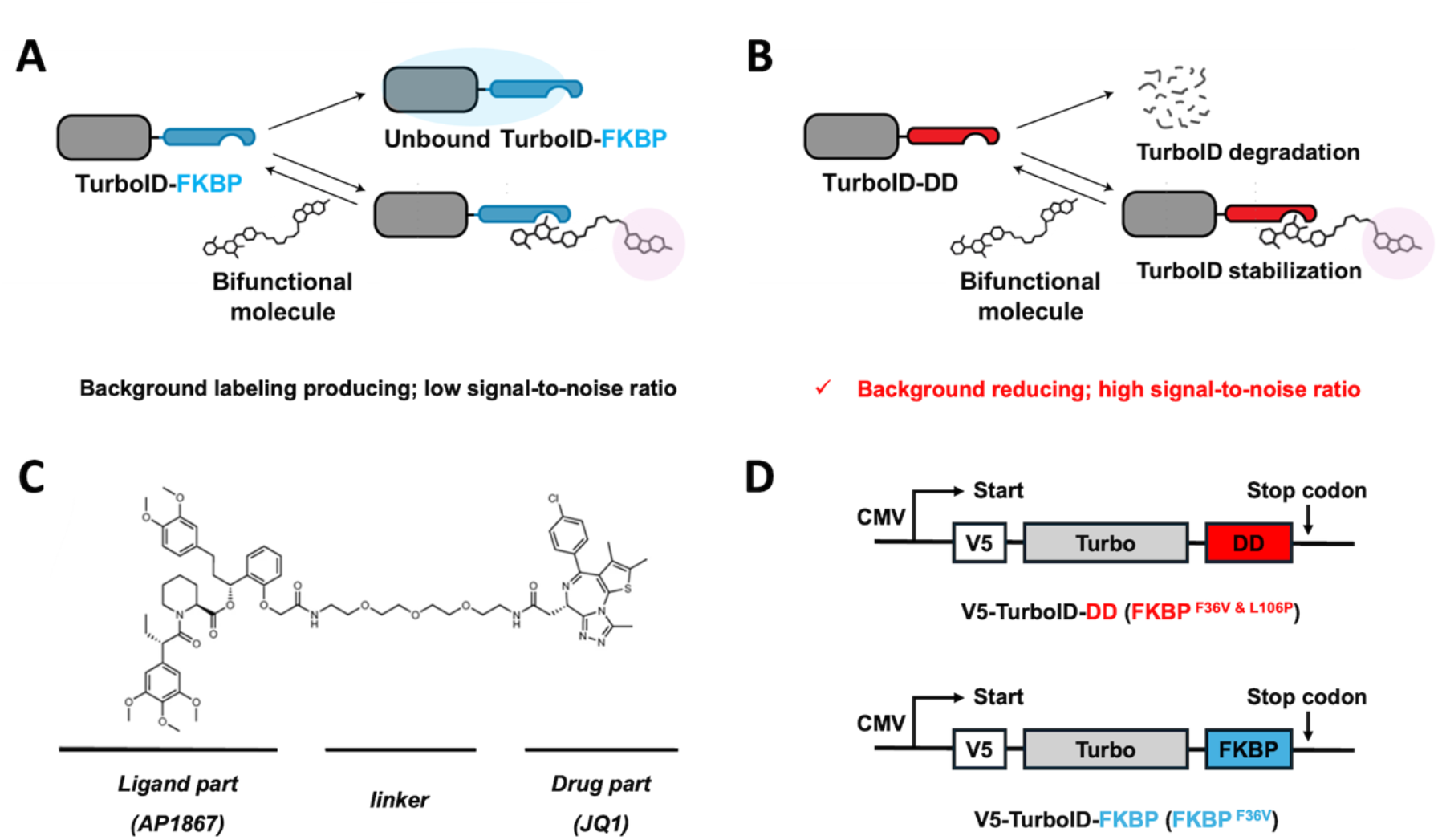
A destabilised TurboID system for drug target identification and interactome mapping. **(A)** Schematic depicting currently the used TurboID-FKBP in targetID and BioTAC. The fusion protein is active even in the absence of the targeting small molecule. **(B)** Schematic depicting a TurboID protein fused to destabilized FKBP version, DD. The fusion protein is degraded in the absence of a bifunctional molecule, leading to reduced background labelling. **(C)** Chemical structure of bifunctional molecule used in this study, NICE-01 with annotations of key components. **(D)** The expression constructs of V5-TurboID-DD/FKBP with position of mutations indicated in parentheses.

To check whether NICE-01 stabilises TurboID-DD as has been observed for DD-tagged YFP ^[11]^, we tested various concentrations of NICE-01 and different exposure times for the cells in the culture medium (Figure 2). A TurboID-FKBP protein, which served as a control, was detected at similar levels, regardless of the NICE-01 concentration or incubation time (Figure 2A and 2B, right panels). In contrast, the amount of TurboID-DD increased with the concentration of NICE-01 and incubation time, reaching a peak after an 8-hour incubation at a concentration of 2.5 µM (Figure 2A and 2B, left panels). We observed no toxic effect of NICE-01 at a concentration range from 0.25 - 5 µM (Supporting Figure S1), consistent with observations made for HCT116 colon cancer and U2OS osteosarcoma cell lines ^[13]^. To demonstrate that the lower levels of TurboID-DD protein observed derive from its destabilisation, we compared protein levels in HeLa cells treated with NICE-01 and MG132, a proteasome inhibitor ^[11]^. Similar increases in TurboID-DD levels were observed in cells treated with 2.5 µM NICE-01 or 50 µM (or more) of MG132 (Supporting Figure S2), suggesting that the increase in protein levels is due to NICE-01-dependent stabilisation rather than an increase in gene expression. From these experiments, we conclude that TurboID-DD is less stable than TurboID-FKBP in the absence of NICE-01, but its expression reaches similar levels to TurboID-FKBP when NICE-01 is present.

**Figure 2.**
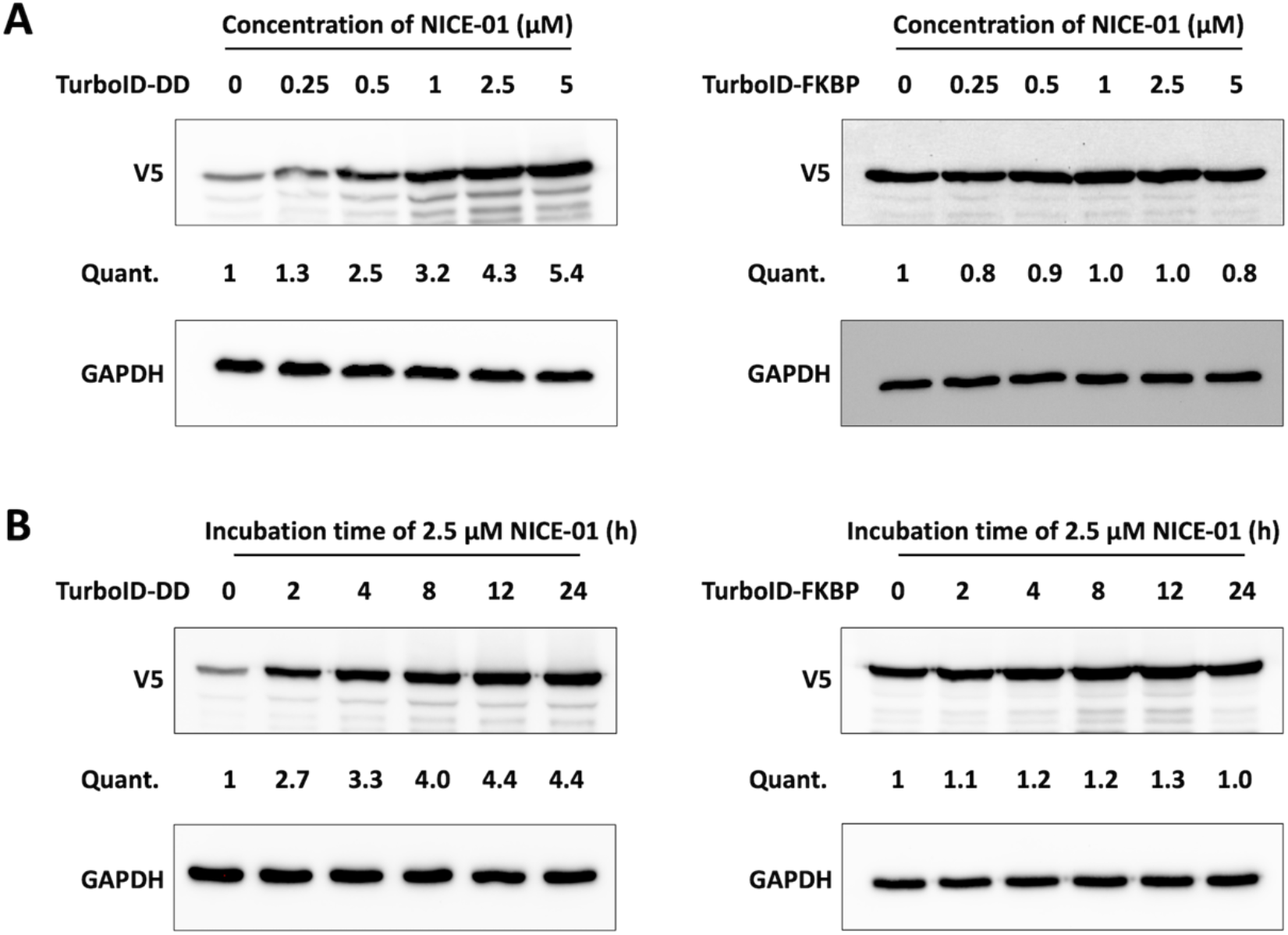
NICE-01 stabilizes TurboID-DD in a dose-dependent manner. **(A)** Immunoblot analysis of V5-tagged TurboID-DD (left) and TurboID-FKBP (right) after exposure of cells to various concentrations of NICE-01 for 24 h. GAPDH is used for normalization of quantification. **(B)** Immunoblot analysis of V5-tagged TurboID-DD (left) and TurboID-FKBP (right) after exposure of cells to 2.5 µM NICE-01 for indicated time. GAPDH is used for normalization of quantification.

### A destabilized TurboID shows improved biotinylation specificity in BioTAC

Next, we tested if the observed destabilization seen for TurboID-DD also results in changes in biotinylation of proteins. Whereas no biotinylation besides background was observed in untransfected HeLa cells, TurboID-DD or TurboID-FKBP expression resulted in a marked increase of biotinylation (Figure 3A, bottom). However, increase in overall biotinylation was less strong in TurboID-DD expressing cells in the absence than in the presence of NICE-01.

**Figure 3.**
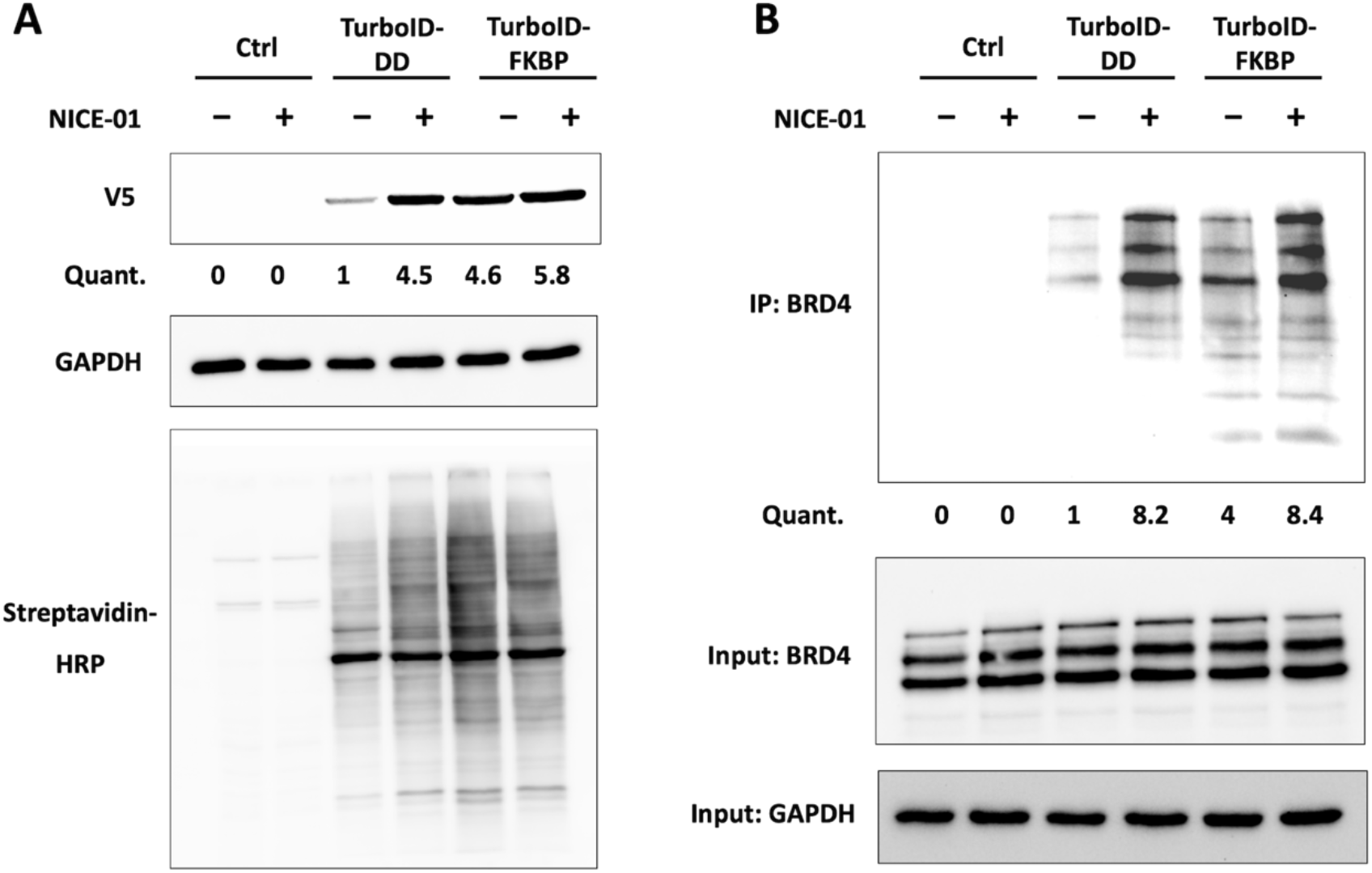
Specific biotinylation by TurboID-DD but not TurboID-FKBP depends on stabilisation by NICE-01. **(A)** Top: Expression of V5-tagged TurboID-DD and TurboID-FKBP in the absence and presence of 2.5 µM NICE-01. GAPDH is used for normalization of quantification. Bottom: Detection of biotinylated proteins by streptavidin-coupled HRP. **(B)** Top: Immunoblot analysis for biotinylated BRD4 after streptavidin affinity purification from cell lysates of untransfected Hela cells (Ctrl) or Hela cells transiently expressing the TurboID-DD/FKBP protein. All cells were treated with 2.5 µM NICE-01 for 24 h. Bottom: Immunoblot analysis of input samples for the presence of BRD4 and GAPDH.

This contrasts with cells expressing TurboID-FKBP where NICE-01 had little influence on biotinylation by TurboID. These differences in the biotinylation intensities reflect the TurboID protein levels as detected by the V5 tag (Figure 3A, top). In support of this, a NICE-01 dependent accumulation of TurboID-DD can also be detected *in situ* by immunofluorescence against the V5 tag of the fusion protein (Supporting Figure S3). Whereas the signal representing TurboID-FKBP is independent of the presence of NICE-01, the intensity of the signal showing TurboID-DD is much stronger in cells treated with NICE-01 than in those without. In addition, we observed a translocation of both proteins into the nucleus upon NICE-01 treatment. This has been observed before ^[8]^ and likely reflects a NICE-01 dependent binding of TurboID-DD and TurboID-FKBP to its targets in the nucleus.

Next, we wanted to test if the specific biotinylation of a known target protein of NICE-01 is improved by TurboID-DD versus TurboID-FKBP. BRD4 was chosen since it is a well-defined target of the JQ1 part of NICE-01 ^[12]^ and has been used in the original BioTAC approach ^[6b]^. Based on the naming of the original approach as BioTAC we named our modified version ddBioTAC (destruction domain assisted biotin targeting chimera). After proximity labeling, biotinylated proteins were isolated by streptavidin affinity purification and analyzed by western blot for the presence of BRD4 (Figure 3B). No biotinylated BRD4 was isolated from extracts from control cells but from cells expressing TurboID-DD or TurboID-FKBP. Importantly, biotinylation of BRD4 by TurboID-DD in the absence of the targeting molecule NICE-01 was fourfold reduced compared to TurboID-FKBP whereas a similar increase in biotinylation was observed in the presence of the molecule. As a net result, we observe a more than eightfold increase in enrichment of biotinylated BRD4 in TurboID-DD expressing cells +/-NICE-01 compared to a twofold enrichment in TurboID-FKBP expressing cells. This indicates a fourfold reduction of biotinylation in cases where the proximity labeling enzyme is not directed to its target.

To expand our analysis to proteins other than BRD4 itself, we combined proximity labeling with steptavidin-based pulldown and mass spectrometric analysis. Three independent replicates for TurboID-FKBP and TurboID-DD in the presence of NICE-01 were analyzed by bottom-up proteomics. In addition, we included three replicates of control experiments with biotinylation performed in the absence of NICE-01 (‘DMSO control’). Biotinylation with biotin, quenching, lysis and capturing was essentially done according to previously published protocols ^[14]^ (see Supporting Information). The captured proteins were further measured by liquid chromatography-tandem mass spectrometry (LC-MS) after in-gel tryptic digestion. Downstream data processing was performed as label free quantification (LFQ; see Supporting Information). With this approach we observed an excellent correlation between the replicates (Supporting Figure S4). We next evaluated the enrichment of identified proteins in the presence and absence of NICE-01 (Figure 4).

**Figure 4.**
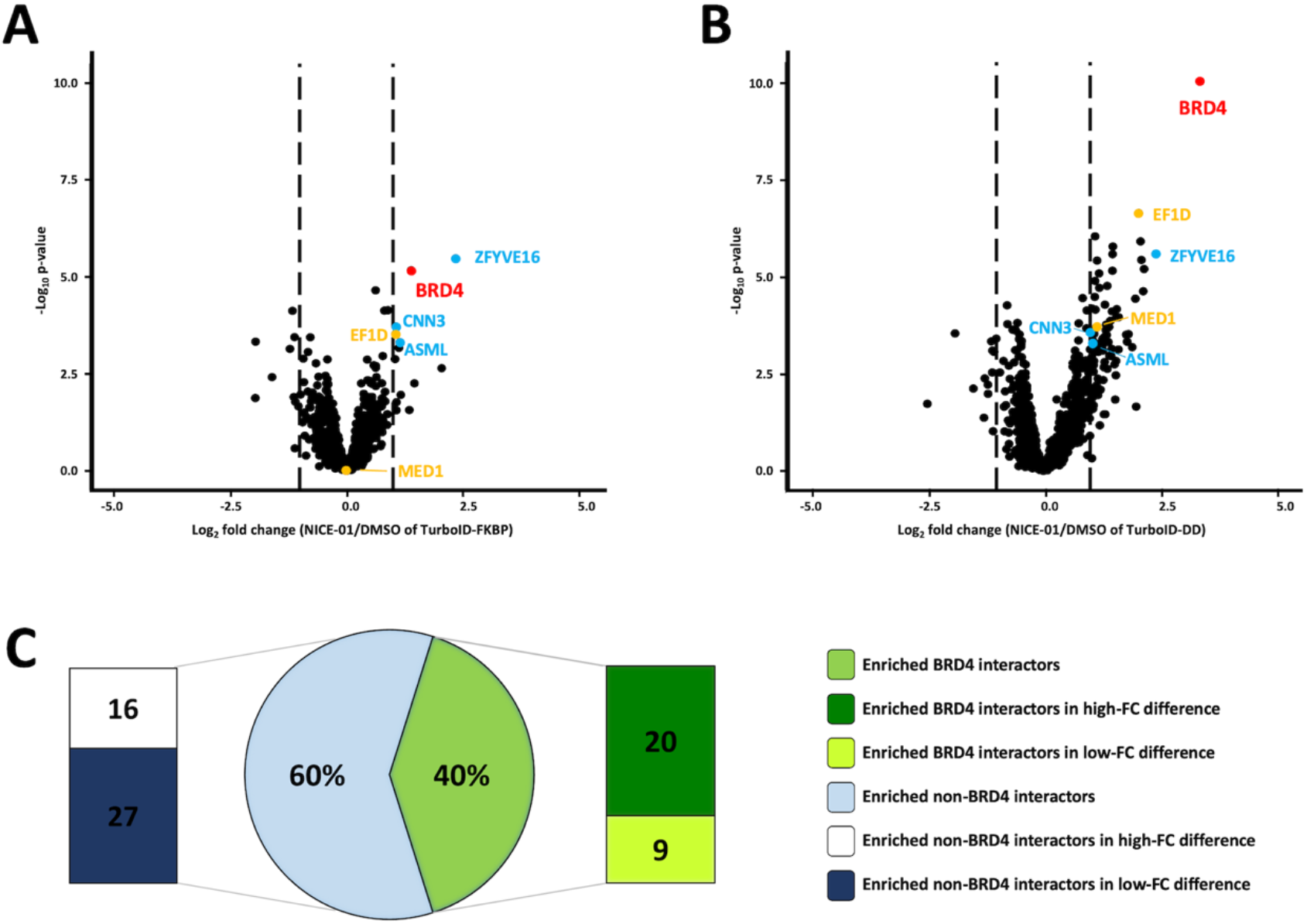
NICE-01-stabilized V5-TurboID enzyme enables accurate identification of drug target and reliable assessment of drug interactome in a high signal-to-noise ratio manner. **(A)** Volcano plot displaying relative fold change of streptavidin-enriched protein abundance following treatment of Hela cells transiently transfected with TurboID-FKBP either DMSO or 2.5 μM NICE-01 for 24 h and 500 μM biotin for 30 min at 37 °C. **(B)** Volcano plot displaying relative fold change of streptavidin-enriched protein abundance following treatment of Hela cells transiently transfected with TurboID-DD either DMSO or 2.5 μM NICE-01 for 24 h and 500 μM biotin for 30 min at 37 °C. Red indicates direct target, Orange indicates reprensentive literature-curated BRD4 interactors from BioGRID. Blue indicates reprensentive non-BRD4 interactors according to BioGRID. Only proteins with valid intensity values in both conditions are depicted, all enriched proteins are listed in Supporting Table S1. All samples run as n=3 replicates. **(C)** Pie chart of BRD interactor categorization. All 72 enriched proteins except BRD4 in TurboID-DD were classed into high/low fold change difference between TurboID-FKBP and –DD. Numbers of BRD4 interactors or non-BRD4 interactor according to BioGRID in high/low fold change difference groups were labelled for comparison.

Compared to TurboID-FKBP cells, >10fold more proteins were selectively enriched by TurboID-DD dependent biotinylation (Figure 4), using at a fold change (FC) cutoff of log_2_FC=1 and p-value of 0.05. Importantly, the relative enrichment of the drug target BRD4 itself was significantly higher in cells expressing TurboID-DD in the presence of NICE-01 (red label, Figure 4A and 4B).

BRD4, as the mainly direct target of JQ-1, stood out more prominently from all proteins enriched by TurboID-DD biotinylation compared to TurboID-FKBP. However, no other bromodomain proteins, including BRD2 and BRD3 were found to be enriched. A potential reason for this is the high ambiguity of interpreting mass spec results regarding BRD2/3/4 due to their similarities in sequence, especially in drugID related setups ^[15]^. However, due to the higher affinity of JQ-1 for BRD4 than BRD2 and BRD3, we interpret our mass spectrometry results as BRD4 ^[12]^. Nonetheless, our results regarding BRD4 suggest that TurboID-DD can better differentiate a true drug binder from non-binders than TurboID-FKBP. In addition, two BRD4 interactors, MED1 ^[16]^ and EF1D ^[17]^ are significantly more enriched in the TurboID-DD dataset after NICE-01 treatment than in the TurboID-FKBP dataset. We measured a twofold enrichment for both proteins. Importantly, the relative enrichment of proteins that are not directly interacting with BRD4 (e.g CNN3, ASML, or ZFYVE16) does not change between TurboID-FKBP and TurboID-DD, indicating that ddBioTAC can better discriminate than BioTAC between drug targets and their direct binding partners versus non-interactors. In total, we could identify 73 proteins (including BRD4) as significantly enriched in the proxisome of TurboID-DD (Supporting Table S1). However, only 40% of these proteins (n= 29) are labeled as potential BRD4 interactors (Figure 4C) according to BioGRID ^[18]^; https://thebiogrid.org/117036/summary/homo-sapiens/brd4.html). To investigate this further, we ranked the 72 identified proteins. Their enrichment over the no-drug control was compared to that seen in cells expressing TurboID-FKBP, creating a fold change (FC) difference, which was used for ranking (Supporting Table S1). The listed proteins were divided into high-FC and low-FC groups (using a threshold of log2 FC = 0.9). Of the 29 proteins classified as potential BRD4 interactors, 20 showed up in the group with high FC, while 27 of 36 proteins not classified as potential BRD4 interactors are in the low FC group. We conclude from this that ddBioTAC improves the identification not only of the drug targets but may also be better suited than BioTAC for the identification of proteins that are proximal to or interact with the drug target.

## Conclusion

Although we have only tested ddBioTAC with a single drug (JQ1) that targets bromodomain proteins, we can claim that there is a significant improvement in this case compared to the first-generation BioTAC system. Replacing (mini)TurboID with a fusion that is constantly destabilised it in the absence of ortho-AP1867/Shld1 allows expression from even strong constitutive promoters, such as the CMV promoter, without affecting the signal-to-noise ratio between target proteins and those labelled by the untargeted enzyme. While ddBioTAC may have limitations similar to those of BioTAC when applied to ubiquitin ligase recruiting molecules ^[7, 19]^, as well as relatively long labelling times, these can be overcome by the recently established APEXTAC system ^[7]^. Nevertheless, ddBioTAC has certain advantages over APEXTAC. Labelling in the latter system is based on an engineered ascorbate peroxidase (APEX2) and requires hydrogen peroxide (H_2_O_2_). The potential toxicity of H_2_O_2_, its ability to induce or alter stress-sensitive pathways and its limited tissue penetration restrict its use ^[7]^. As these limitations do not apply to biotin ligase-based systems (including TurboID), the presented ddBioTAC will be advantageous as a second-generation TargetID approach for identifying drug targets and target-associated proteins.

## Supporting information

Supporting Information Xue etal

## Supporting Information

Supporting information for this article (Experimental Methods, Supporting Figures S1-S4, and a Supporting Table S1) are available. The authors have cited additional references within the Supporting Information.^[30, 31]^

## Acknowledgements

Proteomic studies were performed at the Proteome Center Tuebingen. The authors would like to thank Mirita Franz and Anke Biedermann for planning and conducting the LC-MSMS analyses. YX was supported by the Chinese Scholarship Council (CSC) -Tübingen PhD Program.

