## Supporting Information Xue etal for "DRUG TARGET IDENTIFICATION VIA A CONDITIONALLY STABILIZED TurboID ENZYME"

**Content**

| | |
|---|---|
| Experimental Methods | 1 - 3 |
| Supporting References | 4 |
| Supporting Figures S1 – S4 | 5 - 7 |
| Supporting Table S1<br>Fold change difference TurboID-DD and TurboID-FKBP | 8 - 9 |

### Experimental Methods

#### Plasmid Construction:

All plasmids were generated using standard cloning procedures via T4 DNA Ligase. Restriction enzymes and ligases were obtained from New England Biolabs (NEB). For the plasmids harboring V5-TurboID-DD, the DD ($FKBP^{F36V\&L106P}$) gene was cloned from pSF4 TetCMV intron 20xGCN4 Renilla FKBP Stop 24xMS2v5 SV40 CTE polyA [1] (#119945, Addgene) and inserted into V5-TurboID-NES_pCDNA3 [2] (#107169, Addgene) for mammalian cell expression. To generate the V5-TurboID-FKBP ($FKBP^{F36V}$) expressing plasmid, the above-mentioned plasmid encoding V5-TurboID-DD was mutated at $Pro^{106}$ position to Leu by site directed mutagenesis. *Escherichia coli* strain TOP10 was used as the transformation host. Plasmids were confirmed via Sanger sequencing before use.

#### Cell Culture:

HeLa (EM2-11ht) cells were cultured in high-glucose DMEM (Gibco, #11965092), supplemented with 10% fetal bovine serum (FBS) (Gibco, #10437028). The cells were maintained in a 37 °C incubator with 5% $CO_2$ in a water saturated atmosphere. For sub-cultivation and seeding, cells were washed with DPBS (Sigma-Aldrich, #D8537), detached with 0.25% trypsin/EDTA (Sigma-Aldrich, #T4049) and resuspended in fresh DMEM/FBS. Sub-cultivation was performed every 3–5 days.

#### Cell Viability Assay

HeLa cells were cultured and plated in a 96 well plate (Greiner, #655180) at 40,000 cells in 100 µL culture media per well, then returned to the incubator overnight. The culture media was aspirated and 100 µL of DMEM with NICE-01 (Tocris Bioscience, #8088) in different concentrations was added to each well. After 24 hours the media was aspirated and a mixture of 10 µL CCK-8 solution (Sigma-Aldrich, #96992) with 100 µL DMEM was added to each well. After incubation for 4 hours at 37 °C, the absorption was measured at OD450 using a plate reader.

#### Biotin Labeling

After transfection for 24 h with Fugene HD (Promega, #E2311), the medium was replaced by warm NICE-01/DMSO containing DMEM medium for the other 24 h incubation. Then cells were changed into warm NICE-01/DMSO and biotin containing DMEM medium to initiate labeling with 500 µM biotin. Cells were incubated at 37 °C for 30 min before stopping the reaction by placing the cells onto ice and washing five times with ice-cold (4°C) DPBS.

#### Streptavidin-based Enrichment of Biotin-Labeled Proteins

For each sample, 100 µL streptavidin magnetic beads (Cytiva, #28985799) were washed twice with 1 mL RIPA lysis buffer. For each sample, beads were incubated with 1 mg protein and an additional 1 mL RIPA lysis buffer. Incubation was done at

4°C overnight or at room temperature for 2 h with rotation. Afterwards beads were collected using a magnetic rack and the supernatants transferred into fresh microcentrifuge tubes. The beads were washed twice with 1 mL RIPA lysis buffer for 2 min at room temperature, followed by a wash with 1 mL 1 M KCl for 2 min at room temperature. This was followed by four further wash steps, once with 0.1 M $Na_2CO_3$ (1 mL, 10 s), once with 2 M urea in 10 mM Tris-HCl pH 8.0 (1 mL, 10 s), and twice with RIPA lysis buffer (1 mL per wash, 2 min at room temperature). The enriched material was eluted from the beads by boiling for 10 min in 30 µL of 3x SDS sample buffer supplemented with 2 mM biotin and 20 mM DTT. Beads were separated from the supernatant with a magnetic rack and the eluates used for immunoblot analysis or mass spectrometry.

#### Immunoblot Analysis and Antibodies

For all western blots, 20–50 µg of protein was separated on SDS-PAGE gels, transferred to nitrocellulose membranes, and stained by Ponceau S (5 min in 0.1% (w/v) Ponceau S in 5% acetic acid/water). The blots were blocked in 5% (w/v) milk in 1x TBST (Tris-buffered saline, 0.1% Tween 20) for at least 30 min at room temperature. Blots were incubated with primary antibodies in 5% milk (w/v) in 1x TBST overnight at 4°C or for 1 h at room temperature, and washed three times with 1x TBST for 10 min each, followed by incubation with secondary antibodies in 3% milk (w/v) in 1x TBST for 1 h at room temperature. The following primary antibodies were used: mouse anti-V5 (1:5000 dilution, Thermo Fisher, #R960-25), mouse anti-GAPDH (1:10000 dilution, Proteintech, #60004-1-Ig), and rabbit anti-BRD4 (1:1000 dilution, Bethyl Laboratories, #A301-985A-M). For the streptavidin-HRP conjugate (1:3000, Thermo Fisher, #S911), the blots were blocked in 3% BSA in 1x TBST overnight at 4°C, then incubated with 0.5 µg/mL streptavidin-HRP conjugate for 1 h at room temperature. The blots were washed three times with 1x TBST for 10 min each before development with Pierce™ ECL Western Blotting substrate (Thermo Fisher, #32209). Images were acquired on a ChemiDoc Imaging System (Bio-Rad).

#### In-gel digest and Mass Spectrometry

Coomassie-stained gel pieces were in-gel digested with trypsin, desalted and peptide mixtures were analyzed on a Vanquish Neo UHPLC coupled to an Orbitrap Exploris 480 mass spectrometer (both Thermo Fisher). Prior to MS-based analysis, peptides were separated on an in-house packed ReproSil-Pur C18-AQ 1.9 µm resin (Dr Maisch GmbH Ltd.) 20 cm analytical column (75 µm ID PicoTip fused silica emitter (New Objective). For gradient elution of the peptides, solvent A (0.1% formic acid) and solvent B (0.1% formic acid in 80% acetonitrile) were used across a 60 min gradient with a flow rate of 200 nL/min at 40 °C. The mass spectrometer was operated in positive ion and in data-dependent acquisition mode. MS and MS/MS spectra were generated at a resolution of 60k. In all measurements, sequenced precursor masses were excluded from further selection for 30 s. The target values for MS/MS fragmentation were $10^5$ charges and, for the MS scan, $3 \times 10^6$ charges. The mass spectrometry proteomics data have been deposited to the ProteomeXchange Consortium via the PRIDE partner repository with the dataset identifier PXD077481.

Generated raw files were further processed with MaxQuant software, version 2.5.0.0. The spectra were searched against a homo sapiens database obtained from Uniprot (downloaded 30th of January 2024; 104,581 entries), and 285 commonly observed contaminants. The data were processed with a setting of 1% for the FDR (False Discovery Rate), i.e. with an estimation that 1% of all identifications are false-positive.

For downstream data analysis, proteins were required to be quantified in three replicates of TurboID-DD and TurboID-FKBP groups. Data was then uploaded and analyzed using *Provision*, a web-based platform for rapid analysis of proteomics data processed by MaxQuant [3]. Label-free quantification (LFQ) intensities were $log_{10}$-transformed prior to statistical analysis. Minimum unique peptides were set as 2. Control imputation metrics were set within a width of 0.3 and a downshift of 1.8. P-values and fold differences (FC) were output by *Provision* and for graphical visualization principal component analysis (PCA) and volcano plots showing statistical significance $-log_{10}$ (*p*-value) versus $log_2$ FC were produced by the platform.

**Immunofluorescence Microscopy**

After transfection for 24 h with Fugene HD, the medium in each well was replaced by warm NICE-01/DMSO containing DMEM medium for another 24 h incubation. Then cells were fixed and penetrated by cold methanol at -20°C for 20 min. Cells were washed three times with ice-cold DPBS and blocked with 1% (wt/vol) BSA in DPBS at 4°C for at least 30 min. After removing the blocking solution, cells were incubated with mouse anti-V5 antibody (1:1000) in 1% (wt/vol) BSA in DPBS at 4°C for 1 h. After three gentle washes with 1% (wt/vol) BSA in DPBS, cells were incubated with Alexa Fluor 488 conjugated goat anti-mouse antibody (1:1000, Thermo Fisher, #A32723TR) in 1% (wt/vol) BSA in DPBS at 4°C for 1 h. Finally, cells were washed three times with 1% (wt/vol) BSA in DPBS and stained with Dapi (200 ng/mL) in 1% (wt/vol) BSA in DPBS at 4°C for 15 min. Coverslips with cells were mounted onto glass slides using a mounting medium and allowed to set overnight at room temperature in the dark. Images were acquired using microscope Zeiss Cell Observer with appropriate filter settings.

### Supporting Figures

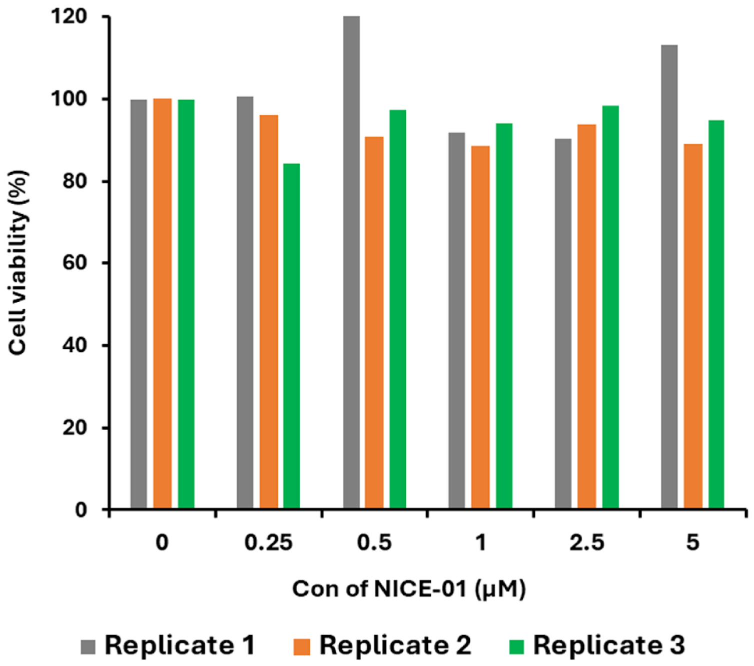

**Figure S1.** CCK-8 viability assay of Hela cells treated with different concentrations of NICE-01 for 24 h (three replicates for each concentration). The cell viability percentage was normalized using a control group and calculated as: cell viability (%) = (Abs (drug, cells) – Abs (no drug, no cells)) / (Abs (no drug, cells) – Abs (no drug, no cells)) * 100%.

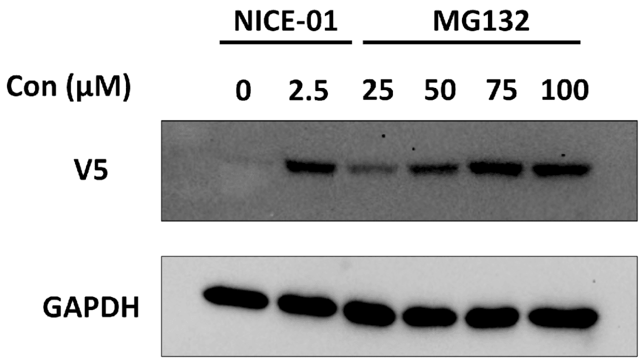

**Figure S2**. The degradation of V5-TurboID-DD is mediated by the ubiquitin-proteasome system. Immunoblot analysis for the indicated proteins from whole cell lysates (WCL) of Hela cells transiently expressing the V5-TurboID-DD cassette treated with the indicated doses of NICE-01 for 24 h or MG132 for 8 h.

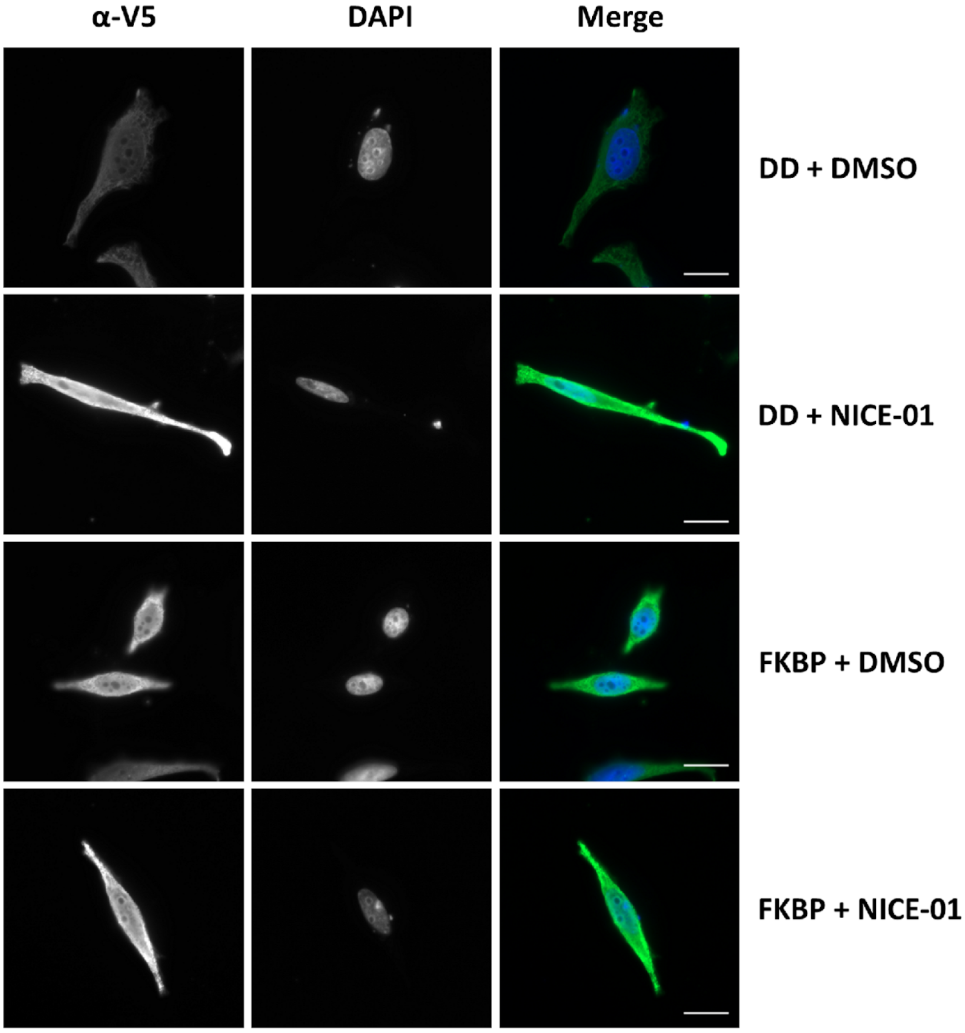

**Figure S3.** Expression and localization of V5-TurboID-DD/FKBP in HeLa cells. Representative fluorescent microscope images of HeLa cells transiently transfected with V5-TurboID-DD/FKBP and treated with DMSO/NICE-01 for 24 h. Left column: V5 (V5-TurboID-DD/FKBP), middle column: DAPI (nucleus), right column: merge of V5 and DAPI signals. Scale bar = 20 μm.

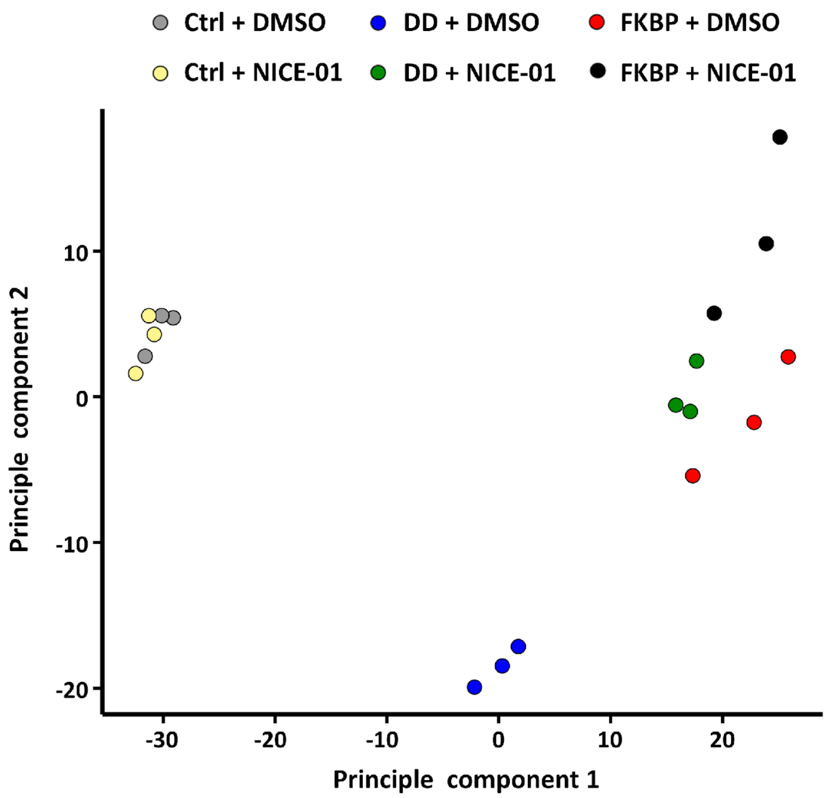

**Figure S4.** Principal component analysis (PCA) representing proteomics data from the comparative analysis of control cells and TurboID-DD/FKBP transfected cells. The PCA plot represents proteins with 3 biological replicates showing distinct proteomic profile differences between DMSO-treated and NICE-01 treated cells, as shown in matching colors.

**Supporting Table S1.** Fold change difference ($log_2$) between TurboID-DD and TurboID-FKBP among all enriched proteins in TurboID-DD group.

| High-FC difference group | | | | Low-FC difference group | | | |
|---|---|---|---|---|---|---|---|
| GeneName | $Log_2$ FC (FKBP) | $Log_2$ FC (DD) | $Log_2$ FC difference | GeneName | $Log_2$ FC (FKBP) | $Log_2$ FC (DD) | $Log_2$ FC difference |
| RBM12 | -1,21137 | 1,401402 | 2,612770123 | CARF | 0,447054 | 1,293287 | 0,846233 |
| MARE1 | -0,18692 | 1,963587 | 2,150502914 | SAHH2 | 0,516796 | 1,362336 | 0,84554 |
| FHOD1 | -0,3248 | 1,816968 | 2,141765278 | CD2AP | 0,247337 | 1,079704 | 0,832367 |
| LYRIC | 0,043413 | 2,131533 | 2,088120162 | ACLY | 0,348434 | 1,17289 | 0,824456 |
| IST1 | 0,054865 | 2,092551 | 2,03768646 | SYTC | 0,266505 | 1,082632 | 0,816127 |
| NUCL | -0,50053 | 1,300764 | 1,801298779 | MTRR | 0,658554 | 1,465553 | 0,806999 |
| E3W994 | 0,017136 | 1,787984 | 1,770847267 | TWF2 | 0,505283 | 1,299632 | 0,794349 |
| CAPR1 | -0,30631 | 1,38804 | 1,694353845 | RBGP1 | 0,788648 | 1,565689 | 0,777041 |
| S23IP | -0,06744 | 1,473712 | 1,541148603 | HP1B3 | 0,831264 | 1,605704 | 0,77444 |
| ZC3HF | 0,704399 | 2,149803 | 1,445403889 | UBP15 | 0,684572 | 1,458078 | 0,773507 |
| NPM | -0,33058 | 1,101855 | 1,432436545 | SWP70 | 0,877514 | 1,60218 | 0,724666 |
| BCAR1 | -0,1244 | 1,266787 | 1,391185785 | SRPRA | 0,353857 | 1,074362 | 0,720505 |
| HTSF1 | 0,090073 | 1,477527 | 1,387453383 | VIGLN | 0,328235 | 1,045448 | 0,717213 |
| COR1C | 0,188427 | 1,547612 | 1,359185471 | MADD | 0,745983 | 1,446515 | 0,700532 |
| PP1G | -0,11327 | 1,216974 | 1,330240629 | KLC1 | 0,645294 | 1,343095 | 0,697801 |
| RECQ1 | 0,238386 | 1,552047 | 1,31366052 | IF2P | 0,426218 | 1,118377 | 0,692159 |
| LAP2A | -0,19704 | 1,095946 | 1,292982494 | LYPA2 | 0,72751 | 1,412511 | 0,685002 |
| SAFB1 | -0,19367 | 1,058327 | 1,251999461 | TLN1 | 0,454265 | 1,103196 | 0,648931 |
| MYPT1 | 0,358161 | 1,558484 | 1,200323249 | CPIN1 | 0,46701 | 1,111213 | 0,644203 |
| SEP11 | 0,082757 | 1,277287 | 1,194530198 | WAC2A | 0,465023 | 1,090963 | 0,62594 |
| TIF1B | 0,07733 | 1,270763 | 1,193433048 | FAS | 0,569821 | 1,188964 | 0,619142 |
| ANXA1 | 0,167494 | 1,349242 | 1,181747268 | FERM2 | 0,457216 | 1,065139 | 0,607923 |
| SC22B | 0,235123 | 1,366308 | 1,131184762 | CAPZB | 0,547413 | 1,092377 | 0,544964 |
| IGBP1 | 0,766828 | 1,896667 | 1,129838998 | ILKAP | 0,51514 | 1,045446 | 0,530306 |
| PUR6 | 0,185737 | 1,314033 | 1,128295933 | LIMC1 | 0,542194 | 1,068317 | 0,526123 |
| MED1 | 0,004484 | 1,13076 | 1,126276474 | NOSIP | 0,828098 | 1,334206 | 0,506107 |
| SERB1 | 0,395574 | 1,430516 | 1,034942043 | VATA | 0,776777 | 1,259208 | 0,482431 |
| PLIN3 | 0,120144 | 1,150957 | 1,030813499 | EP15R | 0,630991 | 1,095724 | 0,464734 |
| G3BP1 | 0,132095 | 1,146013 | 1,013917413 | FA50A | 1,073671 | 1,51917 | 0,445499 |
| EZRI | 0,405009 | 1,389539 | 0,984530295 | A0A096LP25 | 0,691196 | 1,135857 | 0,444661 |
| IF5 | 0,786502 | 1,768972 | 0,982469348 | SYAP1 | 0,779537 | 1,194531 | 0,414994 |
| EF1D | 1,058837 | 2,034183 | 0,975346464 | PP6R3 | 0,644757 | 1,037402 | 0,392645 |
| GLCNE | 1,126597 | 2,074337 | 0,947739734 | JIP4 | 0,760075 | 1,057247 | 0,297172 |
| WDR70 | 0,548438 | 1,483266 | 0,934827571 | ZFY16 | 2,343175 | 2,398869 | 0,055694 |
| SAHH | 0,376319 | 1,308375 | 0,932056342 | CNN3 | 1,062271 | 1,037384 | -0,02489 |
| NIBA2 | 0,245532 | 1,164367 | 0,918835039 | ASML | 1,148352 | 1,033837 | -0,11451 |
| BRD4 | 1,389521 | 3,337651 | 1,948130142 | | | | |

The listed proteins were divided into high-FC and low-FC groups using a threshold of log2 FC = 0.9. Each group contained 36 proteins. BRD4 (blue, at bottom of table) was excluded from the analysis. To investigate BRD4 interactors, listed proteins were sorted in the BioGRID database [4] (https://thebiogrid.org/117036/summary/homo-sapiens/brd4.html). BRD4 interactors according to BioGRID are marked in yellow. Non-interactors of BRD4 interactors are not colour labeled.
